# *In vivo* detection of HIV-1 antisense transcripts in untreated and ART-treated individuals

**DOI:** 10.1101/2024.12.06.627170

**Authors:** Adam A. Capoferri, Rachel Sklutuis, Toluleke O. Famuyiwa, Sachi Pathak, Rui Li, Jason W. Rausch, Brian T. Luke, Rebecca Hoh, Steven G. Deeks, John W. Mellors, John M. Coffin, Jennifer L. Groebner, Fabio Romerio, Mary F. Kearney

## Abstract

Natural antisense transcripts are expressed in eukaryotes, prokaryotes, and viruses and can possess regulatory functions at the transcriptional and/or post-transcriptional levels. *In vitro* studies have shown that HIV-1 antisense transcripts (AST) promote viral latency through epigenetic silencing of the proviral 5′ long terminal repeat (LTR). However, expression of HIV-1 AST *in vivo* have not been convincingly demonstrated. Here, we used single RNA template amplification, detection, and sequencing to demonstrate expression of AST in unstimulated PBMC collected from people with HIV-1 (PWH). We found that AST had high genetic diversity that matched proviruses in cells from blood and lymph nodes. We measured a median of 26 copies of AST per 100 infected cells in PWH on ART and a median of 2 copies per 100 infected cells in PWH not on ART. The expression of HIV-1 AST *in vivo* is consistent with a potential regulatory role in regulation of HIV-1 expression.

---

Antisense transcripts (AST) are RNA molecules transcribed from the opposite strand of a protein-coding gene that can have protein-coding and/or non-coding activities (*1*). AST have been identified in eukaryotes, prokaryotes, and viruses and have been shown to possess regulatory functions at both the transcriptional and post-transcriptional levels via multiple mechanisms (*2*). AST have previously been documented to be encoded by several viruses that infect eukaryotes including members of the *Herpesviridae* (*e.g.* herpes simplex virus-1 and cytomegalovirus) (*3, 4*) and *Retroviridae* (*5–7*) families. One of the best characterized AST in a viral system is the *Hbz* gene in Human T-cell Leukemia Virus Type 1 (HTLV-1). The interplay between *Hbz* (antisense) and *Tax* (sense) RNA expression and their protein products is thought to modulate the regulation of cellular pathways that promote survival and proliferation of HTLV-1 infected cells, thereby influencing the progression into adult T cell leukemia/lymphoma or HTLV-1-associated myelopathy/tropical spastic paraparesis (*8*).

R.H. Miller first provided evidence of an antisense gene (named *asp*) overlapping the *env* gene in the HIV-1 genome (*9*). Several *in vitro* studies have since demonstrated the presence of a Tat-independent negative sense promoter in the HIV-1 3′ long terminal repeat (LTR) (*5, 10, 11*), which drives the expression of multiple antisense transcripts (*12*) with both protein coding (*13*) and non-coding functions (*14*). The presence of HIV-1 antisense protein (ASP)-specific antibodies in serum (*15*) and cytotoxic CD8+ T-lymphocytes (CTLs) in blood samples (*16*) have been detected from people living with HIV (PWH), thus providing indirect evidence for the expression of ASP *in vivo*. Detection of ASP has been reported in eight chronically infected lymphoid and myeloid cell lines during latent and productive infection (*17*). HIV-1 AST have been shown to inhibit viral replication and to promote and maintain latency in stably expressing CD4+ T cell lines (*18, 19*). Although *in vitro* studies suggest that AST function in viral latency, only a few reports have examined its expression *in vivo*. Zapata *et al.* reported low levels of AST in 3 PWH on antiretroviral therapy (ART) (>2 years with undetectable levels of viremia) using real-time PCR in unstimulated CD4+ T cells (*19*) and Mancarella *et al.* (*20*) detected AST in CD4+ T cells collected from both untreated and ART-treated PWH after *ex vivo* stimulation with anti-CD3/CD28. However, no studies have provided sequence evidence for expression of AST in unstimulated cells collected from PWH. Therefore, we set out to determine if we could measure and genetically characterize HIV-1 AST in unmodified peripheral blood mononuclear cells (PBMC) collected from donors on and off ART, and to compare thier expression levels to that of HIV-1 sense transcripts in the same donors. Investigating expression of HIV-1 AST *in vivo* may contribute to our understanding of HIV-1 persistence and reveal new targets for controlling HIV-1 expression without ART.

## Results

### Participants and samples

To determine the levels of AST during chronic infection, PBMC were collected from 3 PWH on ART (*21*) and 5 PWH who were not on ART (**Table 1**) and who were enrolled either at University of California of San Francisco under the SCOPE trial (clinical trial # NCT00187512) or at University of Pittsburgh under the Optimization of Immunologic and Virologic Assays for HIV Trial (IRB# STUDY20040215). All donors provided written informed consent for the study. For one donor (PID 2669), samples collected at four longitudinal timepoints before and after an ART interruption and re-initiation were available.

**Table 1.**
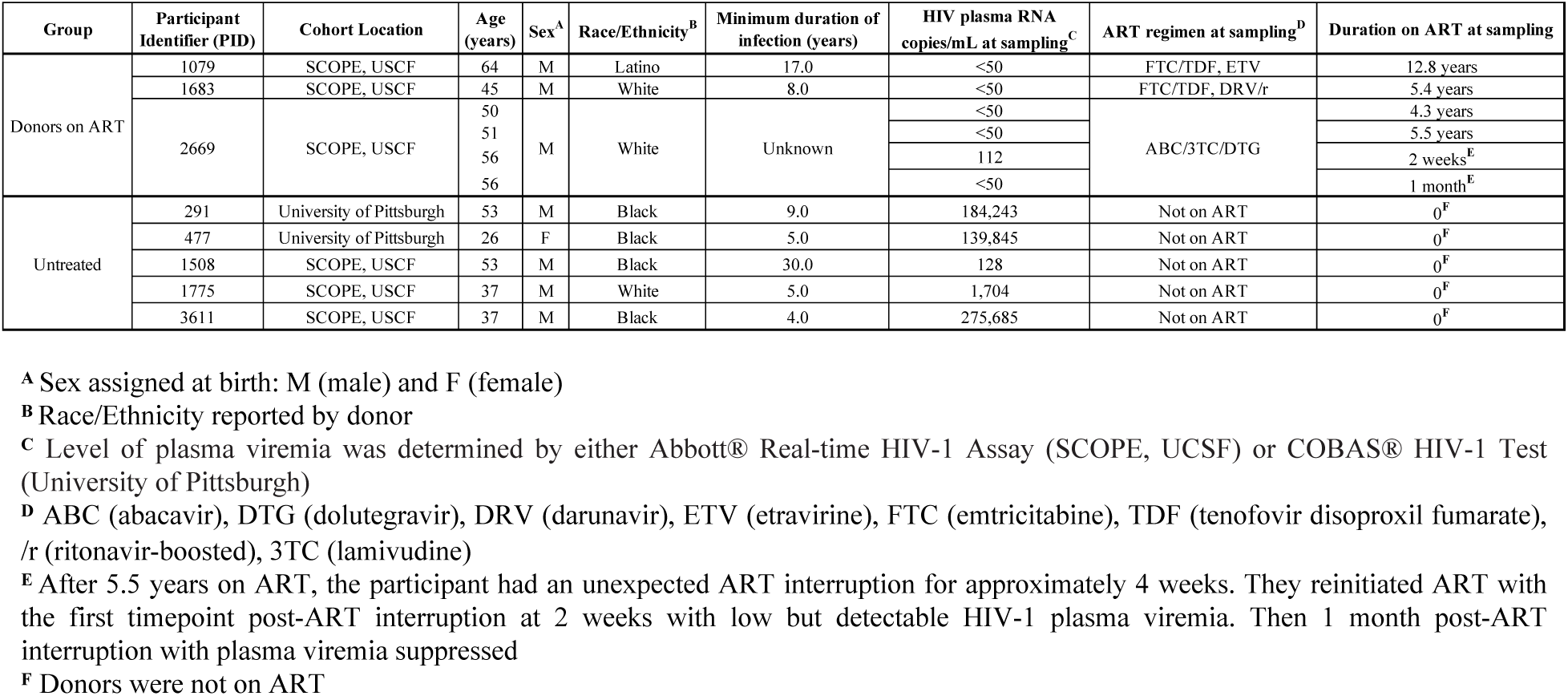
Donor demographics.

Participants were mostly assigned male at birth (n=7/8). Race/ethnicity as reported by donors was Black (n=4/8), White (n=3/8), and Latino (n=1/8). Donors were diagnosed with HIV-1 subtype B with a minimum duration of infection median of 8 years [IQR 5-13 years] prior to sample collection, however this information was unknown for one participant (PID 2669), who was on study more than 5 years. Donors not on ART (untreated) were either ART naïve or not currently on an ART regimen due to a planned or unplanned interruption, and their plasma viremia levels were detectable (median 139,945 range: 128 - 275,685 HIV-1 RNA copies/mL). All but one of the samples collected from PWH on ART had undetectable levels of viremia (<50 HIV-1 RNA copies/mL) with durations of treatment ranging from 2 weeks to 12.8 years. The one sample with detectable viremia on ART (112 HIV-1 RNA copies/mL) was collected from PID 2669 2 weeks after ART re-initiation after an unplanned 4-week treatment interruption. Levels of plasma viremia were measured as HIV-1 RNA copies/mL by either Abbott® real-time assay or COBAS® HIV-1 test.

### HIV-1 AST were detected in ART-treated and untreated PWH, with higher levels in samples collected during ART

Our methods for detecting and measuring levels of HIV-1 AST were modified from Wiegand *et al.* (*22*), Capoferri *et al.* (*23*), and Zapata *et al.* (*19*) and are described in detail in the **Supplementary Materials**. Briefly, to measure levels of HIV-1 AST *in vivo*, we extracted total cell-associated RNA from unmodified donor PBMC with known estimated numbers of HIV-1 infected cells (*24*), synthesized cDNA using participant-specific exogenous oligo-tagged gene-specific primers targeting AST in the *env* coding region (*19*), and quantified the number of AST in each sample with participant-specific anti-*env* primers and probes (primer/probe sequences in **Tables S1-S5**) in a digital PCR format. Prior to testing the donor PBMC, we optimized the AST assay on antisense RNA in the ACH-2 cell line and found these cells to express a median of 41 AST per 100 infected ACH-2 cells (**Supplementary Materials**). Extensive controls were performed to ensure complete degradation of HIV-1 DNA (*22, 23*) and to determine the cut-off for HIV-1 AST (details in **Supplemental Materials**). Equal numbers of no reverse transcriptase wells (negative controls) and experimental wells were included on each PCR plate to ensure no HIV-1 DNA contamination.

AST were detected in all 8 donors (**Fig. 1A & Table S6**). The level of AST in ART-treated samples was a median of 26 copies/100 infected PBMC [IQR 16-47] and the level in untreated samples was a median of ≤ 2 copies/100 infected PBMC [IQR 1-19] (p=0.05, exact p=0.048, Mann-Whitney U-test). Since PID 2669 was overrepresented in the treated group, we also performed the comparison by aggregating the AST in the 2 timepoints prior to the ART interruption, to include only one data point per donor. The re-analysis resulted in a median of 17 AST/100 infected PBMC in the donors on ART vs. ≤ 2 AST/100 infected PBMC in the untreated donors (p=0.21, Mann-Whitney U-test). In PID 3611 (untreated), only 1 AST molecule was detected in 244 infected cells, which was below the cut-off for the assay (details for cut-off in **Supplementary Materials**), indicating that the AST in this donor sample was ≤ 0.4 copies/100 infected PBMC. In PID 1775 (untreated), detection of 33 copies/100 infected PBMC was an outlier among the other untreated PWH (≤ 0.4-5 copies/100 infected PBMC), as confirmed by the Grubbs’ Test for Outliers (G=1.8, α=0.05) (*25*). In PID 2669 (treated), AST were detected in samples collected at all 4 timepoints: 4.3 and 5.5 years after ART initiation and 2 weeks and 4 weeks after ART re-initiation following a brief treatment interruption. Samples after ART re-initiation had higher levels of AST than samples collected prior to ART interruption (49% of 133 infected PBMC vs. 25% of 155 infected PBMC) (p=3.4x10^-9^, binomial test).

**Fig. 1.**
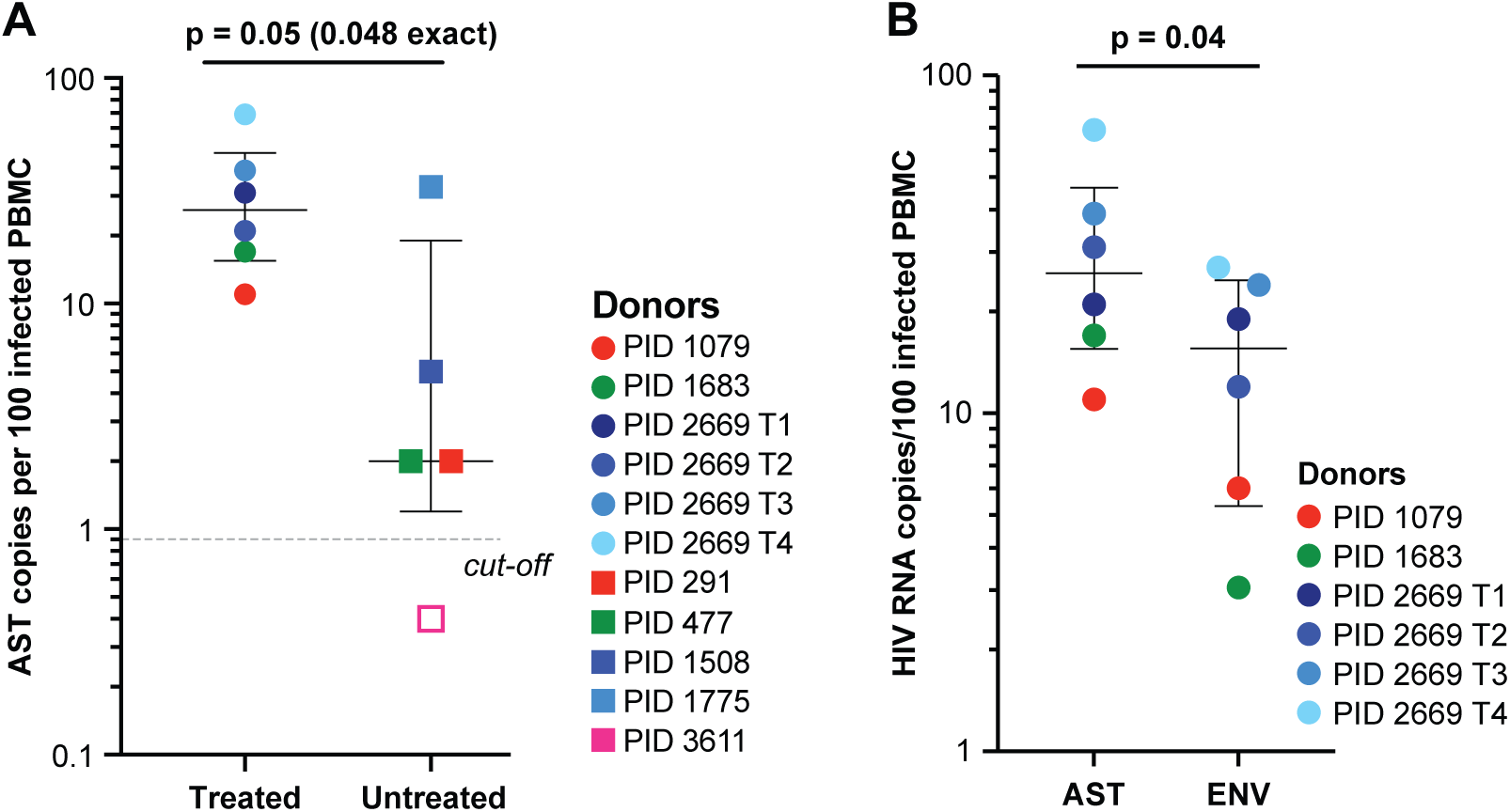
Levels of HIV-1 AST and sense *env* transcripts in PWH. (**A**) Levels of HIV-1 AST in ART-treated and untreated PWH. PWH on ART (circles) and PWH not on ART (squares), each color represents an individual, and PID 2669 has multiple shades of one color to indicate longitudinal sampling. P value determined with Mann-Whitney U-test. Open shape indicates a sample where AST was detected but was below the assay cut-off. The cut-off was determined as the number of positive RT-wells per number of infected cells assayed across 16 replicates of 100 ACH-2 in a background of 10^5^ CEM cells (see **Supplementary Materials**). (**B**) Detection of AST vs. sense *env* in ART-treated PWH. P value determined with paired t-test.

### HIV-1 AST were detected at modestly higher levels than HIV-1 sense env transcripts in samples on ART

In the samples on ART, we also measured the levels of sense *env* RNA targeting the same sub-genomic region targeted for AST (**Fig. 1B**). Levels of sense *env* RNA (aggregate of unspliced and partially spliced) were quantified similarly as above but as a separate reaction from HIV-1 AST. Levels of AST were modestly higher than sense *env* RNA (median 26 copies/100 infected PBMC [IQR 16-47] vs. median 16 copies/100 infected PBMC [IQR 5-25]) (paired t-test t(5)=2.79, p=0.04).

### Long fragments of HIV-1 AST were amplified from single infected cells from donors on ART

Because of the higher levels of AST in the donors on ART, we were able to amplify longer fragments in this subset. We amplified 1.7-kb fragments of AST spanning from the negative sense promoter in the 3′ LTR to *env* (referred to here as “long AST”). Using a modified version of the CARD-SGS assay (*22, 23*), we used the sequence data to estimate the fraction of infected cells with “long AST” and the levels of AST in single infected cells (**Table S2**; details in **Supplementary Materials**). We found that a median of 4.1% [IQR 1.6 – 5.2%] of infected PBMC had detectable levels of “long AST” with a median of 1.1 copies/cell [IQR 1.0-1.7] at a given point in time, indicating that the frequency of detection of “long AST” was lower than the short AST fragments detected by the digital PCR approach (above). In PID 2669, we found no significant difference in the fraction of cells with “long AST” or the levels of AST in single infected cells across the four time points (Kruskal-Wallis test, H(3)=3.02, p=0.39). Additionally, we found no significant difference when we aggregated the samples prior to ART interruption and after ART-reinitiation (p=0.83, Mann-Whitney U-test).

### Phylogenetic analysis of AST reveals that their expression originates from a diverse population of proviruses

Having successfully amplified the 1.7-kb segments of HIV-1 AST in the donors on ART by RT-PCR, we sequenced the resulting PCR products and performed phylogenetic analyses to compare the genetics of the AST to the proviruses in the same population of infected cells (**Fig. 2**). Standard HIV-1 *env* DNA single-genome sequencing using PBMC from all 3 donors and lymph node mononuclear cells (LNMC) from 2 of the donors was performed previously (*21*) and the data used here as a reference. Neighbor-joining trees were reconstructed using HIV-1 *env* DNA from PBMC and LNMC and ∼1.0-kb of the AST in the same genetic region. Trees were rooted on the HIV-1 subtype B consensus sequence. Symbols on the trees show HIV-1 *env* DNA from PBMC (black triangles), HIV-1 *env* DNA from LNMC (blue triangles), and AST (multicolored squares where each color is obtained from a different aliquot of PBMC). AST matching HIV-1 *env* DNA in PBMC or LNMC are indicated with black arrows; AST from infected probable T cell clones are indicated with blue arrows. In PID 1079, a cell expressing high levels of AST is indicated with a red arrow. The genetic diversity of AST was measured by average pairwise distance (APD) with predicted hypermutant sequences removed (*26*).

**Fig. 2.**
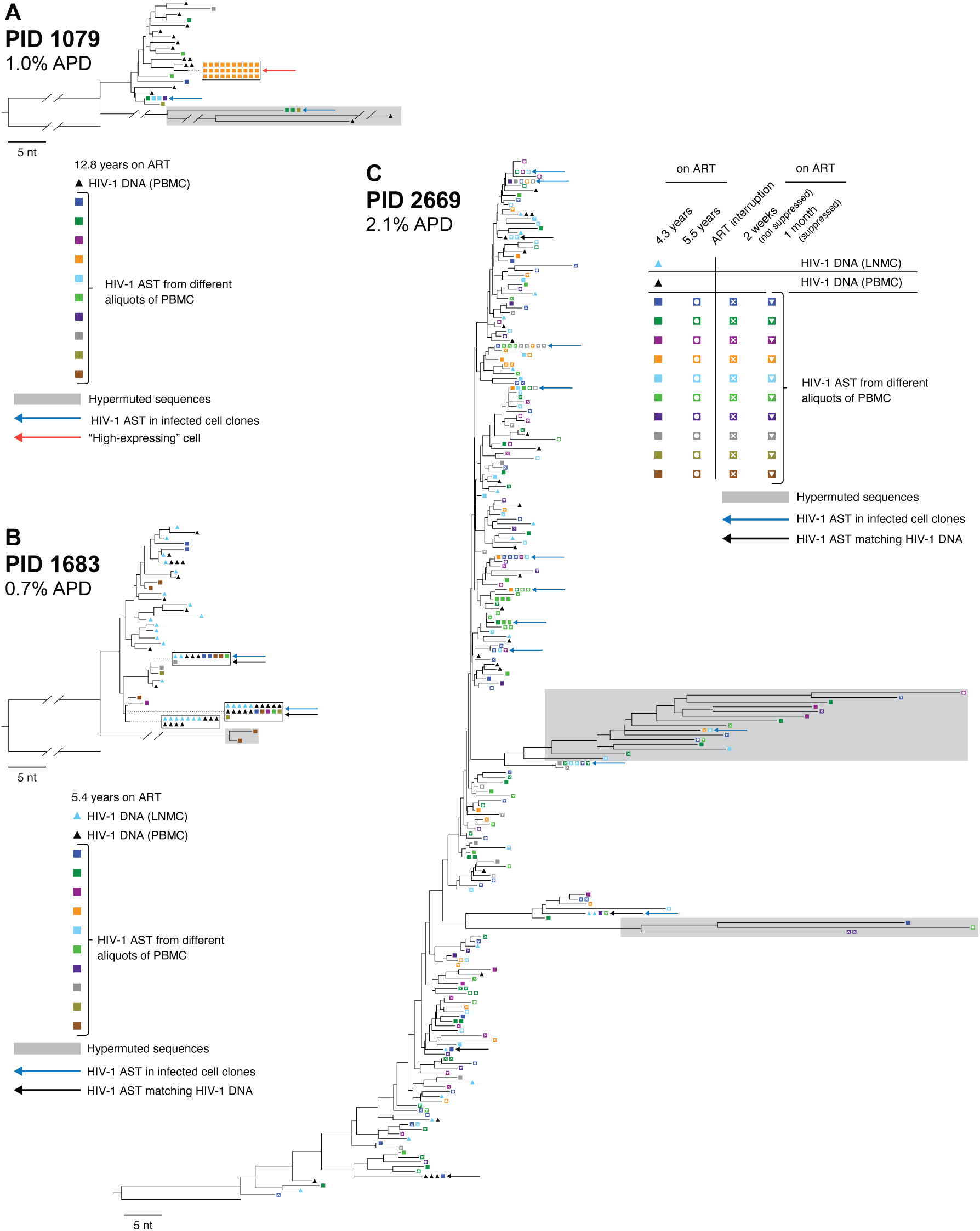
Distance trees of AST in donors on ART. HIV-1 AST sequenced using the modified CARD-SGS assay (**Supplementary Materials**). Percent average pairwise distance (APD) was calculated without hypermutants. HIV-1 AST sequences were aligned with the HIV-1 DNA *env* sequences (previously reported in McManus *et al*. (*21*)). Black triangles show proviruses from PBMC, light blue triangles show proviruses from lymph node mononuclear cells (LNMC), and squares are the intracellular AST from PBMC with each color representing a different aliquot of PBMC. Blue arrows indicate identical AST sequences found in >1 aliquot of PBMC, black arrows indicate identical AST sequences that matched either PBMC or LMNC HIV-1 DNA, and the red arrow indicates a “high AST-expressing” cell. Predicted hypermutant sequences are shaded in gray. Trees are rooted on consensus subtype B *env*.

**Fig. 3.**
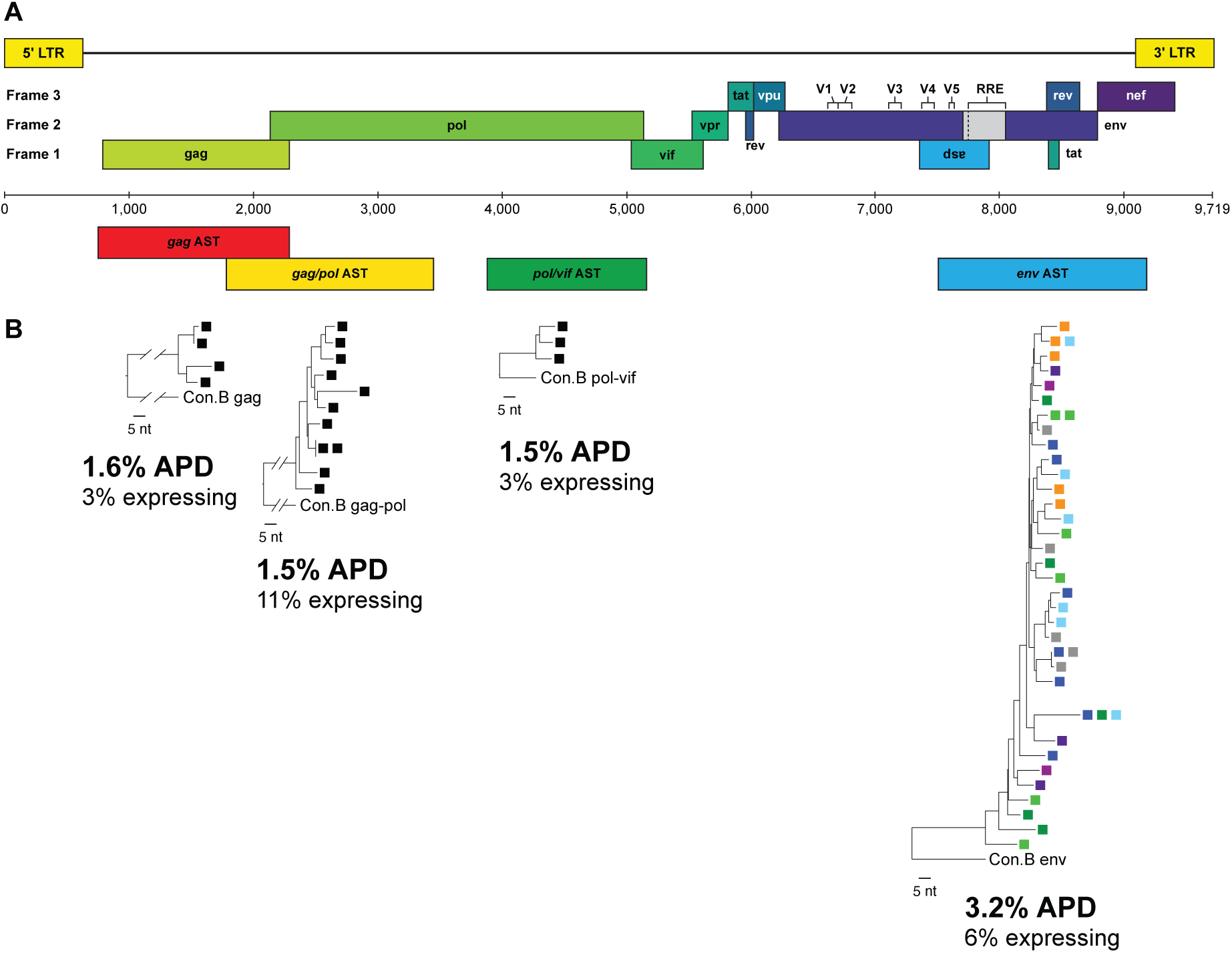
Detection of antisense transcripts along the proviral genome. (**A**) HIV-1 genome map indicating sub-genomic regions for AST amplification. (**B**) Modified AST single-genome sequencing using exogenous oligo-tagged primers to synthesize cDNA and to PCR amplify antisense transcripts along the genome. RNA from aliquots of 90 infected PBMC were extracted from PID 2669 Timepoint #4 for each sub-genomic region (anti-*gag*, *gag/pol*, *pol/vif*) and denoted by a single colored square. p-distance neighbor-joining phylogenetic reconstruction of positive amplicons were generated and percent APD and fraction of expressing cells are reported. The AST tree for anti-*env* is extracted from Fig. 2C, where up to 10 aliquots of PBMC were used. Each color square represents a different aliquot of PBMC from Timepoint #4.

We detected a high genetic diversity of AST (ranging from 0.7–2.5%) matching a diverse population of proviruses in both PBMC and in LNMC (1.1-2.1%), indicating that a wide variety of proviruses can express AST (**Fig. 2**). In one aliquot of PBMC from PID 1079, we found a rake of 30 identical AST, suggesting that they may have originated from the same infected cell (red arrow, **Fig. 2A**). Further, we found identical AST across multiple aliquots of infected PBMC in all 3 donors, indicating that these AST may have originated from infected T cell clones (blue arrows). In PID 1683, we also found identical PBMC AST matching proviruses in both PBMC and LNMC (**Fig. 2B**). While we did not directly assess the presence of AST in LNMC due to limited sampling with fine needle aspirates, these data suggest that AST may be expressed in cell clones that are present in tissues as well as in blood. AST were highly diverse in all 4 samples collected from PID 2669: T1 (2.9% APD), T2 (2.4%), T3 (2.6%), and T4 (2.4%). We also identified 11 likely T cell clones that contained some cells with AST (blue arrows). Eight of the 11 clones were found to persist across multiple timepoints, mostly either before or after the ART interruption, and only very rarely persisting both before and after the interruption, suggesting that ART interruption may influence the populations of T cells that express HIV-1 AST (**Fig. 2C**).

### AST can be detected in gag, pol, and env coding regions in vivo

*In vitro* studies have detected AST of varying lengths, including “full-length” (*i.e.*, from the U3/R of the 3′ LTR to *gag*) (*5, 11, 12*) (**Fig. S1A, Supplementary Materials**). We asked if AST could be detected, not only in *env*, but in other HIV genomic regions *in vivo*. We measured AST in *gag* (HXB2: 764-2,281) (*27*), *gag*/*pol* (HXB2: 1,849-3,500) (*22*) and *pol*/*vif* (HXB2: 3,996-5,270) in one donor sample (PID 2669 Timepoint #4) (*28*) (**Fig. S3A**). Cell-associated RNA from three aliquots of ∼90 infected PBMC was extracted and cDNA targeting the antisense strand of *gag*, *gag/pol*, and *pol/vif* was synthesized using exogenous oligo-tagged gene-specific primers, followed by endpoint PCR amplification and Sanger sequencing (primers in **Table S1,4**). We detected AST in all genomic regions assayed. However, we found lower levels of genetic diversity for AST in *gag*, *gag/pol*, and *pol/vif* regions compared to AST in *env* at the same time points (1.5% vs 3.2%) (**Fig. S3B**). Comparable to levels of “long AST” in *env* (1.7kb described above), levels of AST in *gag*, *gag/pol*, and *pol/vif* were 1.0, 1.1, and 1.0 copies/cell, respectively. Although the sub-genomic regions cannot be genetically linked, these findings suggests that AST may span the entire HIV-1 genome in some infected cells.

## Discussion

In this study, we used established quantitative PCR methods (*19*), extensive positive and negative controls, and sequencing to demonstrate the expression of HIV-1 antisense RNA in people living with HIV who are either untreated or are on long or short-term ART. Prior to our study, HIV-1 antisense RNA had been shown to promote and maintain viral latency in stably expressing cell lines by recruiting the enhancer of zeste homolog 2 (EZH2), a core component of the polycomb repressive complex 2 (PRC2), to the HIV-1 5′ LTR (*18, 19*). Recruitment of EZH2 catalyzes trimethylation of lysine 27 on histone H3 (H3K27me3), a suppressive epigenetic mark that promotes nucleosome assembly and suppression of viral transcription. HIV-1 AST are inefficiently polyadenylated and predominately retained in the nucleus to act as a lncRNA (*29*). Although HIV-1 AST have been shown to inhibit viral replication and promote the establishment and maintenance of latency *in vitro* (*19, 30*), studies investigating AST *in vivo* have been limited. Therefore, we sought to determine if AST are expressed *in vivo* in untreated and/or ART-treated PWH, to quantify their levels in bulk and in single infected cells, and to characterize their genetics. To achieve this, we used a digital PCR assay for the detection and quantification of AST in unstimulated PBMC and we used a modified version of CARD-SGS (*22, 23*) to measure the fraction of infected cells with longer fragments of AST (1.7kb from 3′ LTR to *env*), the levels of AST in single infected cells, and the genetic diversity of AST in the unstimulated PBMC populations. Although methods for the CARD-SGS assay have been published previously (*22, 23*), we included detailed methods in the **Supplementary Materials**.

We detected HIV-1 AST in all 8 donors independent of treatment status. However, we found 13-fold higher levels of AST in PBMC samples collected on ART compared to those not on ART, perhaps consistent with AST functioning in the maintenance of viral latency (*19, 30*). An alternative interpretation could be that, in untreated donors, there are higher levels of transcription from the 5′ LTR. RNA polymerase collision (*i.e.*, transcriptional interference, either “sitting duck” or “roadblock”) or RNA:RNA hybrid formation between 5′ LTR-driven and 3′ LTR-driven transcription may suppress the expression of AST. Except for one donor on ART (PID 1079) where we identified one cell that may have had about 30 copies of AST, the level of AST in single cells was very low, with a mean of 1.2 copy/cell (range 1-30 copies/cell). The low levels of HIV-1 AST are consistent with those reported for other eukaryotic AST. In about 25% of protein-coding genes, AST are expressed at approximately 1 or a few copies per cell at any given time (*31*). In contrast, some AST can be expressed at very high levels, as observed for human *MALAT1* at about 150 TPM (*32*). This range of antisense lncRNA expression has been well documented across cellular and tissue types in the Functional Annotation of the Mammalian Genome (FANTOM) (*33–35*), the Genotype-Tissue Expression (GTEx) consortium (*32*), the Encyclopedia of DNA Elements (ENCODE) project (*31*), and the Long non-coding RNA Knowledgebase (*36*).

In the 3 donors on ART, we amplified and sequenced the 1.7-kb fragment of HIV-1 AST overlapping *env* from about 80 infected cells per sample. We found high genetic diversity of the AST including matches to proviruses in peripheral blood and lymph nodes. In some instances, the AST matched both PBMC and LNMC proviral DNA, suggesting that at least some cells in infected CD4+ T cell clones can express HIV-1 AST. In the one donor on ART with samples from multiple timepoints (PID 2669), we identified probable T cell clones with detectable levels of AST that persisted over time, and one that matched proviral DNA found in both PBMC and LNMC. Interestingly, although identical AST were found across the timepoints before the ART interruption and across the timepoints after the ART interruption and re-initiation, only rarely were identical AST detected both before and after the interruption, suggesting that the treatment interruption may have influenced the populations of T cell clones expressing HIV-1 AST. Further supporting the influence of ART interruption on AST expression was the observation that levels of AST were significantly elevated in samples collected within weeks after ART re-initiation relative to samples collected on longer-term ART (4-5 years on ART). It is possible that levels of AST in donors after ART re-initiation may be different than in long-term treated individuals or that relatively short-lived latently infected cells express AST at higher levels, thus, the effect of ART interruption on levels of AST should be further investigated. Together these data show that HIV-1 AST expression, although typically at low levels, can be detected in probable infected T cell clones in both blood and tissues.

Previous *in vitro* studies have reported multiple HIV-1 AST species using northern blot and 5′ Rapid Amplification of cDNA Ends (5′ RACE): Class I (10kb) (*12*), II (5.5kb) (*11*), III-iii (3kb) (*12*), and IV-ii (2kb) (*5*) (**Fig. S1A**). Our detection of HIV-1 AST in genes other than in *gag*, *gag/pol*, and *pol/vif* at levels comparable to those we observed in *env*, together with the work by Kobayashi-Ishihara *et al*. (*12*), suggest that we detected numerous Class I transcripts, which are full-length, expressed at low levels, and not typically polyadenylated. Relatedly, it is important to note that neither our digital PCR nor our single-genome sequencing assays targeting *env* is informative as to which Class the identified AST transcripts belong.

There are some limitations of this study to consider. Due to the fragility and sparsity of HIV-1 AST, we almost certainly underestimate their frequency. We cannot rule out that HIV-1 AST are short-lived and may not always be detectable. Similarly, there is a potential loss of AST from inefficient reverse transcription due to the thermal stability of secondary and tertiary RNA structures (although the high temperature denaturing step leading into cDNA synthesis is designed to help mitigate such effects), or during the sodium acetate/ethanol precipitation of the antisense cDNA. These recovery limitations are supported by the controls included in the **Supplementary Materials** demonstrating recoveries of only ∼50% for AST spiked samples. Detecting a single copy of HIV-1 AST naturally presents challenges, particularly in determining if the molecules are due to expression by an HIV-1 or host promoter. Readthrough generation of HIV-1 AST can occur when the provirus integrates opposite to a host gene promoter, allowing transcription into the HIV-1 3′ LTR to produce AST. Previous research identifying the transcription start site of HIV-1 AST *in vitro* used infected cell lines and 5′ RACE (*11, 12*). We attempted 5′ RACE on our donor samples to determine if the HIV-1 AST originated from an HIV-1 or host promoter at a single RNA molecule level, but without success due to limitations in assay sensitivity. More sensitive 5′ RACE technologies are needed to determine the transcription start site of single RNA molecules *in vivo*, including HIV-1 AST. However, our finding that there are differences in levels of AST across different cells of a T cell clone may favor expression from the HIV 3′ LTR promoter.

This study is the first to show, with sequence confirmation, the expression of HIV-1 AST in PWH. Additional studies are needed to confirm our observation of higher levels of AST in PWH on ART vs. PWH who are not on ART, and to determine if HIV-1 AST *in vivo* are driven by viral or host promoters. This study, together with those showing the role of HIV-1 AST in inducing and maintaining HIV-1 latency *in vitro* and *ex vivo* (Li *et al.* Romerio, submitted), highlights a previously underexplored potential determinant of HIV-1 persistence both before and during ART, and may lead to new directions for the development of approaches to controlling HIV-1 viremia without ART.

## Acknowledgements

We would like to thank the donors for participating in this study. We acknowledge our collaborative interactions with the Behavior of HIV in Viral Environments Center (U54AI170855). We thank Teresa Burdette and Ann Wiegand for administrative support.

## Financial Support

This work was supported by intramural NCI funding (ZIA BC 011697) to the HIV Dynamics and Replication Program (MFK) and by the Office of AIDS Research. This work was also supported through an Intramural AIDS Research Fellowship (AAC) from the Office of AIDS Research. Other funders include NCI contract No. 75N91019D00024 to BTL, Leidos Biomedical Research, Inc. subcontract 12XS547 to JWM and 13SX110 to JMC. JMC was a Research Professor of the American Cancer Society and supported in part by Research Grants CA R35 200421 and AI R01 184043. The content is solely the responsibility of the authors and does not necessarily represent the official views of the National Cancer Institute, National Institutes of Health, or the Department of Health and Human Services.

## Author Contributions

AAC, RS, JWR, JMC, JLG, MFK designed the study. AAC, RS, TOF, SP, JLG performed experiments. AAC, RS, TOF, JWR, BTL, JLG, MFK analyzed results. AAC, RS, JLG performed sequence data analyses. SGD and JWM provided patient samples. AAC, RS, TOF, FR, MFK wrote the manuscript. All authors reviewed the manuscript. AAC and RS contributed equally to this work.

## Competing Interest Statement

JWM is a consultant to Gilead Sciences, has received research grants from Gilead Sciences to the University of Pittsburgh, and owns share options in Infectious Disease Connect (co-founder) and Galapagos, NV, unrelated to the current work on HIV. JMC is a member of the Scientific Advisory Board and a Shareholder of ROME Therapeutics, Inc. and Generate Biomedicine, Inc. The remaining authors have no potential conflicts.

## Data and Materials Availability

All sequence data are available on GenBank (in process of submission). Sequences from the donors on ART were previously published by McManus and colleagues and can be found in (*21*).

## Supplementary Materials

Materials and Methods

Figs. S1-S4

Tables S1 to S6

References: 37-40

## Supplementary Materials for

### Materials and Methods

#### Participant cohorts, sample collection, and study approval

PBMC were collected from PWH on ART who were enrolled at the University of California of San Francisco in the SCOPE trial (clinical trial # NCT00187512). PBMC were also collected from PWH who were not currently on ART and were enrolled at the University of Pittsburgh in the Optimization of Immunologic and Virologic Assays for HIV Trial (IRB# STUDY20040215). The studies were approved by the University of California San Francisco Institutional Review Board and the University of Pittsburgh Institutional Review Board. All donors provided written informed consent for HIV-1 quantification and sequencing. PBMC were separated using Ficoll, resuspended in FBS with 10% DMSO, and stored in liquid nitrogen (LN_2_) until testing.

#### Overview of HIV-1 antisense transcript (AST) quantification and sequencing approaches

HIV-1 AST quantification and sequencing assays were performed by modifying our cell-associated RNA and DNA single-genome sequencing assay (CARD-SGS) (*22, 23*) (detailed methods below). Briefly, total nucleic acid was extracted from serially diluted aliquots of PBMC collected from PWH, the DNA was digested, and AST cDNA synthesized using a gene-specific primer in the opposite orientation of the *env* coding region (**Fig. S1B**). The cDNA from each aliquot was spread across 96-well PCR plates for amplification (∼50bp) and probe detection to determine the dilution that yielded <30% positive PCR products, indicating that cDNA was primarily amplified from single molecules. By determining the endpoint dilution, we could quantify the number of AST in each aliquot and determine the fraction of the infected cells with AST by using the number of HIV-1 DNA molecules in replicate aliquots for the denominator. HIV-1 DNA was measured with the integrase cell-associated DNA (iCAD) assay (*24*). The number of HIV-1 DNA molecules is a surrogate for the number of infected cells since previous studies showed that most infected cells carry only a single provirus both before and during ART (*37*). For a subset of the samples, a 1.7-kb fragment of AST was also amplified at an endpoint and Sanger sequenced to assess the genetic diversity of the transcripts.

#### Overview of the development and optimization of AST quantification and sequencing

During the development and optimization of the HIV-1 AST quantification and sequencing assays, we controlled for endogenous priming during cDNA synthesis and contamination of HIV-1 DNA. We also determined the sensitivity and background of the AST assays. An overview of these developmental steps is provided here, but detailed protocols for each set of experiments are provided below.

1. To eliminate endogenous self-priming during AST cDNA synthesis (*19, 38*), we used a 5′ end exogenous oligo-tagged gene specific primer (**Table S1**). The tag generates cDNA carrying the exogenous tag sequence to function as the target for the forward primer during first-round PCR amplification; thus, allowing for specific amplification of cDNA molecules containing the exogenous oligo-tag sequence, rather than endogenous self-primed templates.
2. To ensure complete DNA digestion, we used various amounts of the ACH-2 infected cell line (∼1 provirus/cell), spiked into 1x10^5^ or 1x10^6^ uninfected CEM cells (**Fig. S2**). Nucleic acid was extracted, DNase treated, and “cDNA synthesized” without reverse transcriptase. Nested PCR was performed to determine the maximum number of cells that can be assayed to ensure that HIV DNA digestion was complete. We found complete digestion of viral DNA at 100 infected cells per aliquot, in agreement with prior findings with the CARD-SGS assay where <300 infected cells per aliquot was the upper limit (*22*).
3. To optimize the AST assay, we used *in vitro* transcribed AST generated with a pMiniT-AST vector containing a T7 promoter (**Fig. S3**). The concentration of AST was measured by spectrophotometry and 10^4^ copies were spiked into nucleic acid extracted from 1x10^5^ uninfected CEM cells. The number of positives detected in the assay was compared to the known number of transcripts. The optimized AST detection assay had a sensitivity of ∼50% (average of 5,589 copies detected), which could be a consequence of including the exogenous oligo-tag on the cDNA primer reducing cDNA synthesis efficiency or loss of RNA template from the purification steps of the assay.
4. After optimizing the assay using *in vitro* transcribed AST, we tested the assay on 100 ACH-2 cells spiked into 10^5^ uninfected CEM cells (**Fig. S4**). Using the AST assay (43bp amplicon) at endpoint with probe detection, we found a median of 41 AST molecules per 100 infected ACH-2 cells [IQR 25-69] (n=16 replicates). We detected 13 AST /100 infected ACH-2 cells when amplifying and sequencing a 1.7-kb fragment of anti-*env*. The less frequent detection of the 1.7-kb amplicon is not unexpected compared to the 43bp amplicon.
5. To determine the cut-off for the digital (probe detection at an endpoint) AST assay, we performed replicates of “no reverse-transcriptase (RT)” controls on 100 ACH-2 cells spiked into 10^5^ uninfected CEM cells. We found 4 potential false positive signals in 650 infected cells assayed when RT was not included in the cDNA reaction, making our assay cut-off 0.6 AST/100 infected cells. We also tested 5x10^5^ uninfected CEM cells spread across 182 PCR wells and detected 1 false positive, making the false-positive assay background <0.0002/100 cells assayed. To overcome the background, we limited each digital assay to <100 infected cells and included equal numbers of “no RT” controls on each PCR plate.

#### Cell lines, maintenance, and storage

Cell lines used for assay development were uninfected CEM/C1 cells (American Type Culture Collection, #CRL-2265) and HIV-1 infected ACH-2 cells (HIV Reagent Program, #ARP-349). ACH-2 cells are A3.01 cells infected with a single integrated provirus (HIV-1_LAI_) with a defect in TAR (C37T). Complete Medium was prepared as RPMI 1640 supplemented with 1% Penicillin-Streptomycin-Glutamine (Gibco, #10378016) and 10% heat-inactivated fetal bovine serum (HI-FBS). Cell Freeze Medium was prepared as RPMI 1640 supplemented with 10% HI-FBS (CEM/C1 cells) or 10-40% HI-FBS (ACH-2 cells), and 10% DMSO.

Cells were maintained in Complete Medium (as described above) and seeded in 2x vented T-25 flasks (1:3 and 2:3 from vial contents) with a total of 6mL in each flask. Cells were incubated overnight at 37°C, 5% CO_2_ with flasks upright. The following day, the medium was aspirated without disturbing the cells and were replenished with warmed Complete Medium. The cells were subcultured and transferred to a 15mL conical tube for centrifugation at 150x*g* for 10 minutes, then media was aspirated. The cells were resuspended in 1mL Complete Medium and counted by hemocytometer at 1:10 and 1:100 dilutions. Viable cells were seeded at 2-4x10^5^ in new T-25 flasks with 6mL of warmed Complete Medium. If needed, cells were expanded in T-75 flasks (seeding at 4-6x10^5^) with medium volume between 10-20mL. The cells were split every 2-3 days as needed.

The cells were prepared in Cell Freeze Medium at a minimum of 1x10^6^ cells/mL by gentle pipetting of cells and aliquoted into labeled cryotubes. They were placed in a freezing container with 100% isopropanol in outer container to freeze cells at -80°C overnight. The following day, the cells were transferred to LN_2_ storage.

#### Sample Preparation

To thaw viably frozen cells, RPMI 1640 was warmed to 37°C and added dropwise to thaw PBMC for nucleic acid extraction. Each viably frozen cell vial was warmed for ∼2 minutes at 37°C prior to RPMI being added dropwise to the vial. Thawed donor PBMC were aliquoted and centrifuged at 500×*g* for 5 minutes. Supernatant was removed and the pelleted cells were used immediately for downstream assays or were frozen on dry ice and stored back in LN_2_ until ready for use.

#### Generation of control RNA

RNA controls were generated to determine the efficiency of the HIV-1 AST cDNA synthesis and PCR amplification. AST (*anti-env*) were amplified from a pUC57-AST plasmid obtained from Fabio Romerio (Department of Molecular and Comparative Pathobiology, The Johns Hopkins University School of Medicine, Baltimore, MD) using M13 Fwd and M13 Rev primers (**Table S2**). Using Platinum II Taq (ThermoFisher, #14966025), 50μL PCR reactions were carried out with 50ng of plasmid DNA and 10μL of 5X Platinum II PCR Buffer, 1μL of 10mM dNTPs, 1μL of each 10μM primer, 0.4μL of Platinum II Taq polymerase, and molecular-grade water to bring to 50μL. PCR cycling was performed as follows: 94°C for 2 minutes and 45 cycles of 94°C for 15 seconds, 60°C for 15 seconds, and 68°C for 45 seconds, followed by a final extension at 68°C for 1 minute. The PCR product was then purified using a QIAquick PCR Purification Kit (QIAGEN, #28104). The PCR product was cloned using NEB® PCR Cloning Kit (New England Biolabs, #E1202S) where 80ng of the purified PCR product was ligated into a linearized pMiniT 2.0 vector and transformed into the NEB® stable competent 10-beta *E. coli* following manufacturer’s instructions (New England Biolabs, #C3019H). Transformed colonies were picked, grown overnight according to manufacturer’s instructions, and miniprepped using the QuickLyse Miniprep Kit (QIAGEN, #27406). Plasmids were screened for the insert by selective PCR screening using Cloning Analysis Fwd and Rev primers (**Table S2**) with PCR cycling performed as follows: 94°C for 2 minutes and 45 cycles of 94°C for 15 seconds, 60°C for 15 seconds, and 68°C for 45 seconds, followed by a final extension at 68°C for 1 minute. The plasmid was then Sanger sequenced to confirm orientation and sequence (**Table S2**).

To generate control transcripts, 1μg of the AST clone (mentioned above) was digested by restriction enzyme PmeI (New England Biolabs, #R0560S) for AST (*anti-env*) or ZraI (New England Biolabs, #R0659S) for sense *env* transcripts following manufacturer’s instructions. Digested DNA was then purified using the QIAquick PCR Purification Kit (QIAGEN, #28104) and concentrated using ethanol precipitation (described below). RNA was transcribed using HiScribe T7 Quick High Yield RNA Synthesis Kit (New England Biolabs, #E2050S) for AST or the HiScribe SP6 Quick High Yield RNA Synthesis Kit (New England Biolabs, #E2070S) for sense *env* transcripts. Transcribed RNA was purified using a Monarch RNA Cleanup Kit (New England Biolabs, #T2030) and quantified by Nanodrop spectrophotometer. The RNA was diluted to 1x10^6^ copies/mL with 5mM Tris-HCl pH 8.0 and stored at -80°C for temporary storage and LN_2_ for longer storage to help avoid degradation of RNA.

To assess the completeness of digestion of plasmid DNA, approximately 10^4^ copies of the transcripts were spiked into extracted nucleic acid from 10^5^ CEM cells and DNase I treated (described below). The control transcripts were used for cDNA synthesis using random hexamer primers (ThermoFisher, #SO142) in the following reaction: 2.5μL 10mM dNTPs and 2.5μL 2μM random hexamer primer were added to each 20μL RNA sample. The RNA was denatured at 65°C for 10 minutes, then immediately cooled at -20°C for 1 minute. Next, 25μL of SuperScript III Reverse Transcriptase (Invitrogen, #18080-044) master mix was added to the denatured RNA. Master mix was made by combining 10μL 5X first strand buffer, 1μL 0.1M DTT, 12μL RNase-free water, 1μL 40U/μL RNaseOUT recombinant ribonuclease inhibitor (Invitrogen, #10777-019), and 1μL 200U/μL SuperScript III Reverse Transcriptase. cDNA was synthesized at 25°C for 5 minutes, 55°C for 1 hour, 70°C for 15 minutes, then cooled to 4°C. To remove hybridized RNA, 1μL RNase H (5 units) (New England Biolabs, #M0297S) was added and incubated at 37°C for 20 minutes followed by an enzyme heat-inactivation at 75°C for 10 minutes, and the cDNA was cooled to 4°C and used immediately or stored at -80°C.

#### DNA and RNA Extraction

Total nucleic acid was extracted from endpoint diluted aliquots of HIV-infected cells with AST RNA, determined by performing serial dilutions of PBMC until 96-well PCR plates yielded <30% AST positives. Nucleic acid was extracted by adding 100μL 3M guanidine HCl solution (18.75mL 8M guanidinium HCl, 2.5mL 1M Tris-HCl pH 8.0, 0.5mL 100mM CaCl_2_) and 5μL 20mg/mL Proteinase K to a cell pellet, vortexed, and incubated at 42°C heat block for 1 hour. Addition of 400μL 6M guanidine isothiocyanate solution (25mL 6M guanidinium isothiocyanate, 14mL 1M Tris-HCl pH 8.0, 53μL EDTA pH 8.0) and 6μL 20mg/mL glycogen to lysed cell pellet was vortexed, and incubated at 42°C heat block for 10 minutes. Five hundred microliters of 100% isopropanol was added with mixing, followed by centrifugation at 21,000x*g* for 10 minutes at room temperature to precipitate nucleic acids. Supernatant was removed without disturbing the pellet and washed with 750μL 70% ethanol. Precipitated nucleic acid was air-dried. For DNA, the pellet was resuspended in 100μL 5mM Tris-HCl pH 8.0 and ready for use. For RNA, the DNA was digested by either (i) DNase I (Roche, #04716728001) or (ii) ezDNase (ThermoFisher, #11766051):

i. DNase I: The air-dried extracted nucleic acid was resuspended in 38μL DNase buffer and 2μL of 10units/μL DNase I and incubated for 20 minutes in a 37°C water bath. After incubation, 200μL of 6M GuSCN (Sigma, #50983) was added and mixed well, followed by the addition of 250μL of 100% isopropanol. Sample was vortexed for 10 seconds, then centrifuged at 21,000×*g* for 10 minutes to pellet nucleic acid. Supernatant was removed and pellet was washed with 1mL 70% ethanol, followed by additional centrifugation at 21,000×*g* for 10 minutes. Finally, the DNA-digested RNA pellet was air-dried, resuspended in 20μL of 5 mM Tris-HCl pH 8.0, and used immediately for cDNA synthesis. Alternatively, RNA could be temporarily stored in 70% ethanol at -80°C.
ii. ezDNase: The air-dried extracted nucleic acid was resuspended in 32μL 5mM Tris-HCl pH 8.0. Sample was split in equal volumes to control for successful DNA digestion. To ensure successful DNA digestion, manufacturer’s instructions were followed, with volumes of reagents doubled and maximum incubations used. Briefly, 2μL 10X ezDNase Buffer and 2μL ezDNase were added to each sample and incubated at 37°C for 5 minutes. To inactivate the ezDNase, the mixture was incubated at 55°C for 5 minutes in the presence of 2μL 0.1M DTT. Incubations were performed in a thermocycler. Samples were immediately cooled at -20°C and brought up to room temperature before directly moving into cDNA synthesis.

#### cDNA Synthesis

The cDNA was synthesized by adding 2.5μL 10mM dNTPs and 2.5μL 2μM gene-specific tagged cDNA primer (AST cDNA, **Table S1**) to each sample. The RNA was denatured at 85°C for 10 minutes, then immediately cooled at -20°C for 1 minute. Next, 25μL SuperScript III Reverse Transcriptase (Invitrogen, #18080-044) master mix was added to the denatured RNA. Master mix was 10μL 5X first strand buffer, 0.5μL 0.1M DTT, 13.5μL RNase-free water, 0.5μL 40 U/μL RNaseOUT recombinant ribonuclease inhibitor (Invitrogen, #10777-019), and 1μL 200 U/μL SuperScript III Reverse Transcriptase. cDNA was synthesized at 45°C for 1 hour, then cooled to 4°C. To remove hybridized RNA, 1μL RNase H (5 units) (New England Biolabs, #M0297S) was added and incubated at 37°C for 20 minutes followed by an enzyme heat-inactivation at 75°C for 10 minutes. The cDNA was cooled to 4°C and was transferred to a low bind Eppendorf tube and 0.1 vol 3M Sodium Acetate pH 5.5 (Invitrogen, #AM9740) and 1μL 20mg/mL glycogen (Roche, #1090139300) were added. After mixing, 3 volumes 95% ethanol was added, and the entire mixture was vortexed for 10 seconds and incubated overnight at -20°C. The next day, the precipitated cDNA was centrifuged at 21,000x*g* for 20 minutes at room temperature, and supernatant was removed. The pellet was washed with 600μL 70% ethanol, pulse vortexed, and centrifuged for 15 minutes at 21,000x*g* at room temperature. Supernatant was removed and cDNA pellet was air dried and then resuspended in 20μL or 50μL (for digital PCR quantitation) in 5mM Tris-HCl pH 8.0. The cDNA was then used immediately or stored at -80°C.

#### Quantification of HIV-1 infected cells

The number of HIV infected cells was estimated from the levels of HIV DNA in PBMC. For donors on ART, HIV-1 DNA levels were previously measured using the integrase cell-associated single-copy DNA (iCAD) assay (primers in **Table S3**) (*21, 24*). We also measured HIV-1 DNA levels using a modified primer probe set in the Rev Response Element (RRE) adapted from the Intact Proviral DNA Assay (*39*) (**Table S4**). Digital PCR was performed using Lightcycler 480 Probes Master hot-start reaction mix (Roche, #4707494001) in 20μL volumes per reaction. A 2mL master mix was prepared by combining 1mL 2X Lightcycler 480 Probes Master hot-start reaction mix, 12μL 100μM RRE Fwd primer, 12μL 100μM RRE Rev, 2μL 100μM RRE probe, 774μL RNase-free water, and 200μL diluted DNA (DNA was diluted up to 200μL with 5mM Tris-HCl pH 8.0). Digital PCR was performed on the Roche LightCycler480 at a denaturation temperature of 95⁰C for 10 minutes and 55 cycles (95⁰C for 15 seconds and 60⁰C for 1 minute). Equal numbers of “no RT” controls and experimental wells were included on each plate to ensure complete HIV-1 DNA digestion. Two negative (no template, 5mM Tris-HCl pH 8.0) and two positive (100 ACH-2 in a background 1x10^5^ CEM cells of DNA) controls were also included on each PCR plate. The number of positive wells was used to calculate the number of infected cells per million PBMC in each of the donor samples.

#### Design of gene-specific primers and probes

Due to intra- and inter-host HIV-1 sequence variation, we extracted HIV-1 DNA from PBMC (as described above) and performed PCR at endpoint targeting *env* (primers in **Table S1**) and performed Sanger sequencing (primers in **Table S4**). The designed primers for were used accordingly (**Table S5**, donor-specific mutations are shown in bold red).

#### AST quantification by digital PCR

AST digital PCR was performed using Lightcycler 480 Probes Master hot-start reaction mix (Roche, #4707494001) in 20μL volumes per reaction. A master mix was prepared by combining 2X Lightcycler 480 Probes Master hot-start reaction mix, 100μM Fwd primer, 100μM Rev primer, 100μM AST probe, RNase-free water, and cDNA synthesized as described above. Digital PCR was performed on the Roche LightCycler480 at a denaturation temperature of 95⁰C for 10 minutes and 55 cycles (95⁰C for 15 seconds and 60⁰C for 1 minute). Equal numbers of “no RT” controls and experimental wells were included on each plate to ensure complete HIV-1 DNA digestion. Two negative (no template, 5mM Tris-HCl pH 8.0) and two positive (100 ACH-2 in a background 1x10^5^ CEM cells of DNA) controls were also included on each PCR plate.

#### AST quantification and sequencing

AST quantification and sequencing was modified from the original CARD-SGS assay protocol as previously described (*22, 23*) and was used to sequence and quantify HIV-1 DNA and sense and antisense HIV RNA in *gag, pol,* and *env*. Briefly, cDNA or DNA was diluted to a near endpoint in a volume of 200μL and added to 800μL Platinum II Taq (ThermoFisher Scientific, #14966025) master mix containing 200μL 5X Platinum II PCR Buffer, 20μL 10mM dNTPs, 8μL primers (4μL 50μM Forward and 4μL 50μM Reverse primers, for respective region, **Table S1**), 564μL molecular-grade water, and 8μL Platinum II Taq polymerase. The total volume of cDNA and master mix was spread across a 96-well plate (10μL per well). PCR cycling was performed with respect to sub-genomic region (**Table S1**). Following the first round of PCR, each well was diluted with 50μL 5mM Tris-HCl pH 8.0. For nested PCR, 2μL diluted PCR 1 product was transferred from each well to a 96-well plate containing 8μL Platinum II Taq master mix (described above). Nested PCR was performed with respect to sub-genomic region with respective thermocycling conditions (**Table S1**). Nested PCR product was diluted with 50μL 5mM Tris-HCl pH 8.0. Positive wells were identified by GelRed detection (Biotium, #41003) using UV imaging. Positive PCR reactions were sent for Sanger sequencing (**Table S4**).

The levels of HIV expression in single cells were determined by the number of identical sequences in each aliquot. Because the reverse transcription step is known to introduce errors at a rate of about 10^-4^ per sequenced nucleotide of viral cDNA sequences (*40*), RT-PCR variants that differed by a single nucleotide (‘fuzz’) from a group of 7 or more identical sequences within the same aliquot were, conservatively, considered to belong to the same rake of identical sequences. The R script for “defuzzing” is available at https://github.com/michaelbale/RStuff/blob/master/fPECS_fncs.R.

#### Sequence Analysis

Sequences were aligned with MAFFT v7.450 (FFT-NS-1 200PAM/k=2 algorithm) in Geneious Prime ® 2020.2.4. Minor adjustments were performed manually. P-distance neighbor joining trees were reconstructed using MEGA X and rooted on the consensus B HIV sequence. Population genetic diversity was calculated as average pairwise distance (APD) using MEGA 11. APOBEC3-F/G mediated hypermutation was predicted using the Los Alamos National Laboratory HIV Sequence database tool, Hypermut (https://www.hiv.lanl.gov/content/sequence/HYPERMUT/hypermut.html).

#### Statistical analyses

Mann-Whitney U-test, Kruskal-Wallis test, Spearman’s correlation, Grubbs’ Outlier test, and Shapiro-Wilk test for normality for informed Paired T-test were performed using Prism GraphPad 9.2. Spearman’s correlation was selected due to the random variation in biological systems.

To test if there was a difference in the levels of AST pre- and post-ART interruption we used a binomial test. From the two sample donations pre-interruption, a total of 155 infected PBMC were assayed and 39 infected PBMC were found to contain AST (25%). Post-interruption, 133 infected PBMC were assayed and 65 infected PBMC contained AST (49%). A Binomial distribution was used to show that 65 out of 133 is significantly larger than 39 out of 155.

Statistical tests applied are indicated in the text and/or figures/table legends. Additional statistical analyses were performed using R (version 4.3.1).

---

**Fig. S1.**
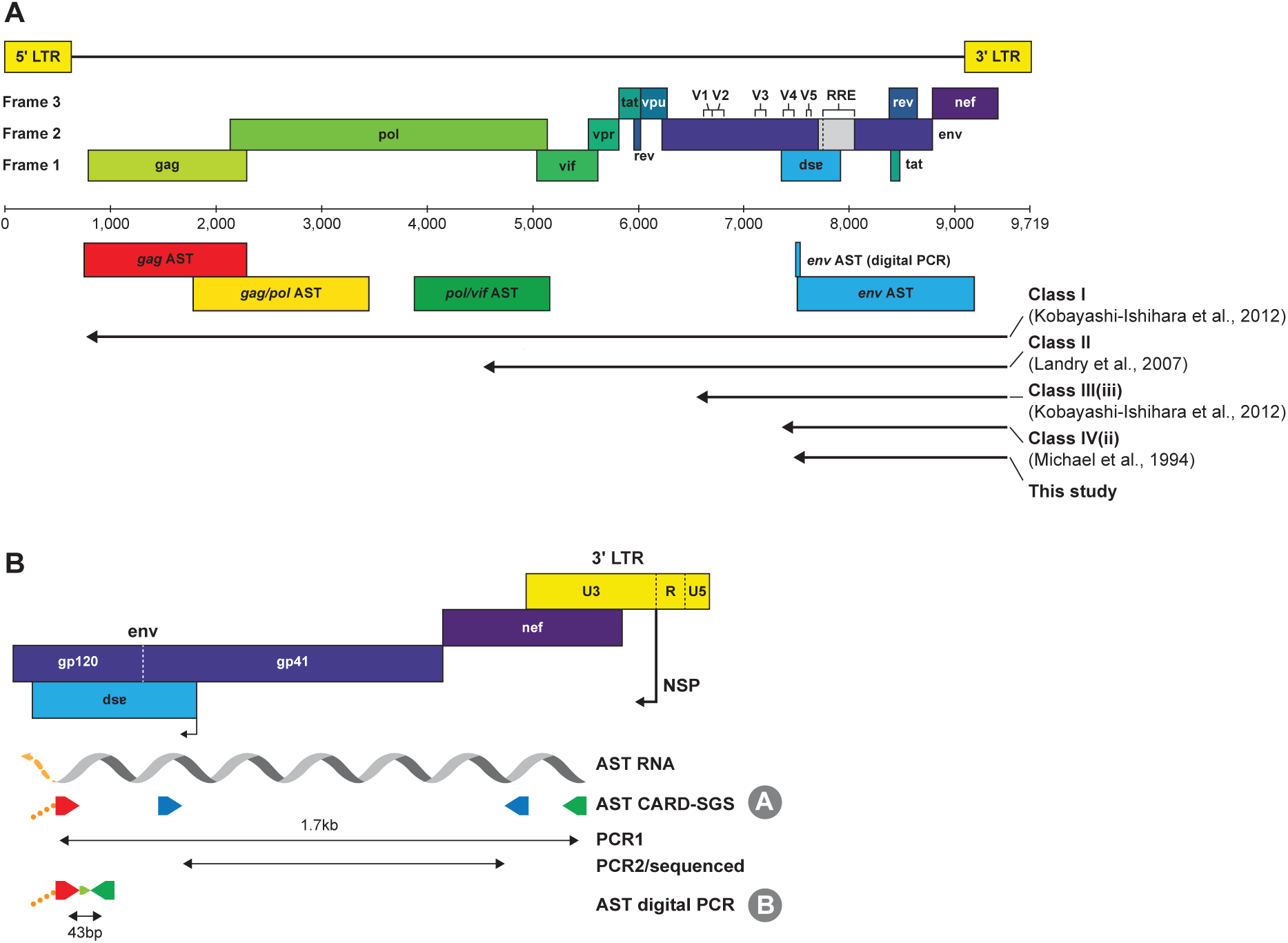
Assays for measuring levels and sequencing AST *in vivo*. (**A**) Measuring AST along the HIV-1 genome. The antisense regions for single-genome sequencing of *gag, gag/pol*, *pol/vif*, and *env* are shown with the anti-*env* location for the digital PCR assay indicated as well. The genome map is relative to HXB2. Different length Classes of AST are defined for Class I (*12*), Class II (*11*), Class III(iii) (*12*), Class IV (*5*), and in our study. (**B**) A detailed view of the locations for the AST CARD-SGS assay (protocol A) and the digital PCR assay (protocol B). The negative sense promoter (nsp) for AST is shown at the 3′ LTR U3/R junction. The orange segment of the AST RNA indicates the potential for the transcript to be longer such as Class I & II. The tag for the primers is bold dashed orange that stems from the cDNA synthesis primer (red). Protocol A for AST CARD-SGS includes PCR1 with the forward primer to the tag to amplify a ∼1.7-kb fragment from gp120 *env* to *nef*/U3, and a nested PCR before sequencing. Protocol B for AST digital PCR, uses the forward primer binding to the tag, and amplifies 43bp. The AST digital PCR assay amplification region overlaps the AST CARD-SGS assay region in *env*.

**Fig. S2.**
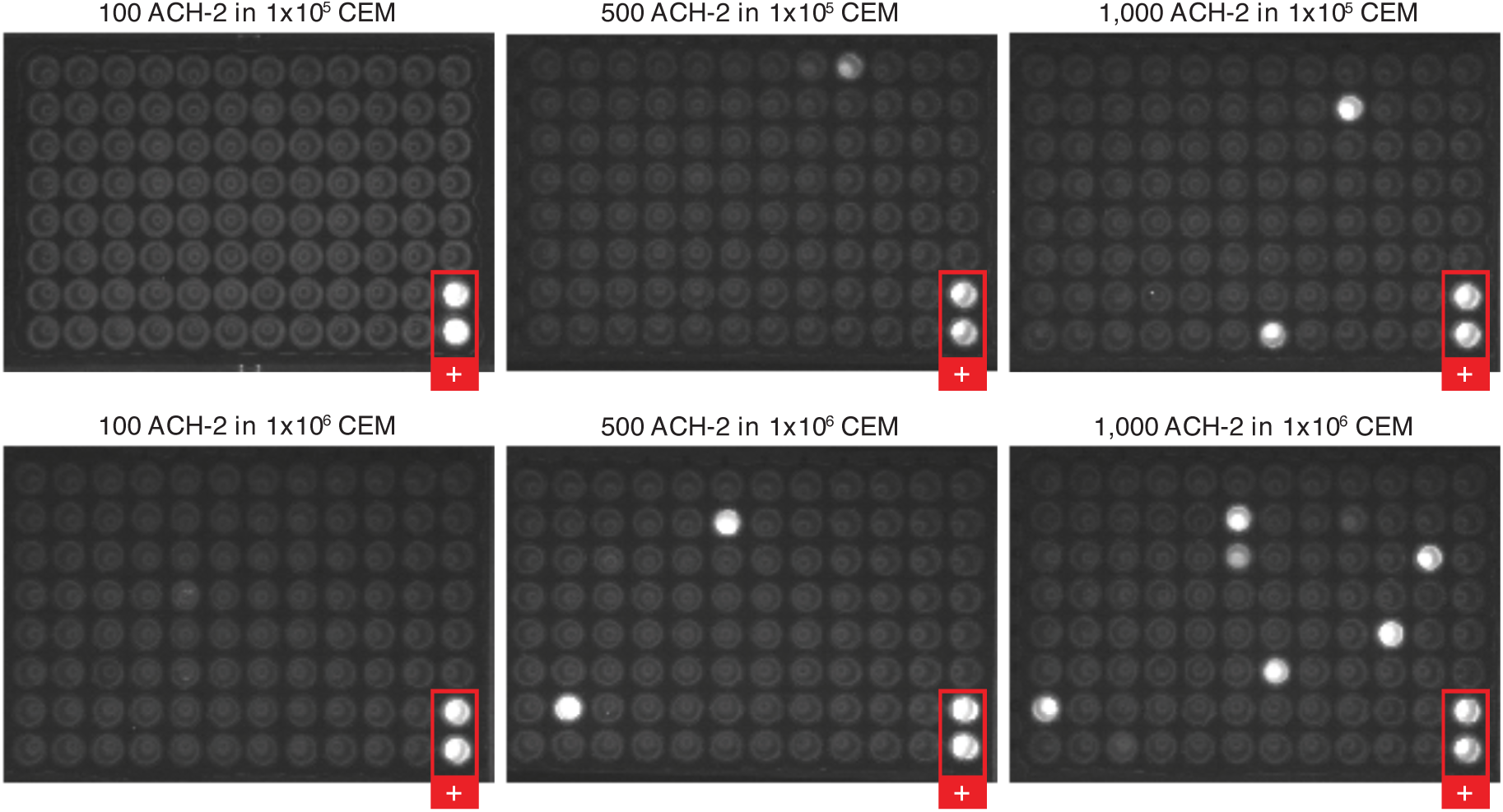
Development of the Antisense Transcripts Cell-Associated RNA and DNA Single Genome Sequencing (AST CARD-SGS) assay – control for DNA digestion. cDNA reactions were performed with the exogenous oligo-tagged RT primer but excluding the reverse transcriptase enzyme, SuperScript III. The reaction was then spread across a 96-well plate and nested PCR was completed. Each plate included positive control wells (red box) that contained ACH-2 DNA.

**Fig. S3.**
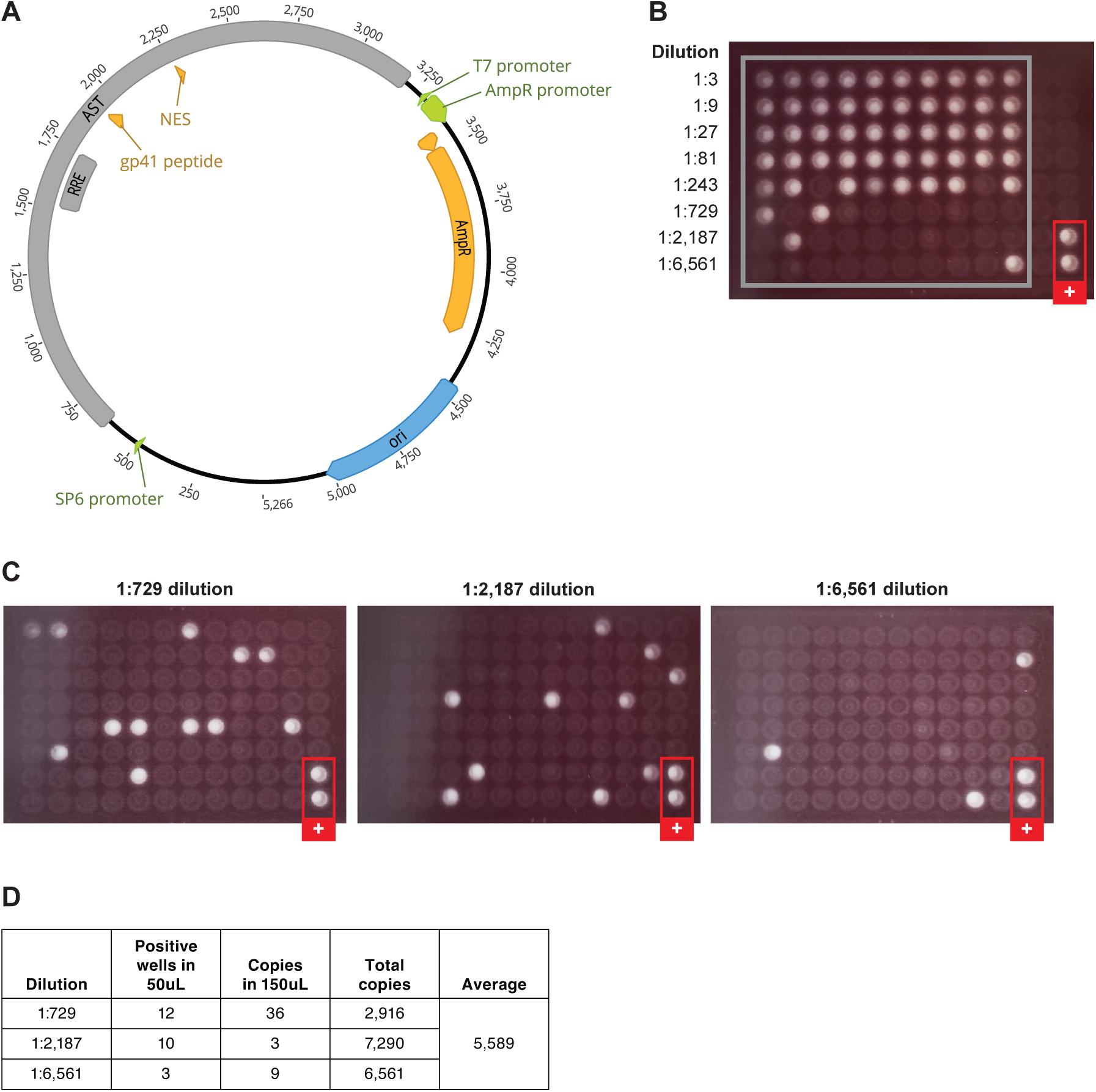
Determining the sensitivity of the AST CARD-SGS. (**A**) Plasmid used for *in vitro* expression of AST (pMiniT Expression Vector). (**B**) PCR replicates of serially diluted *in vitro* transcribed, then reverse transcribed AST for quantification and endpoint determination (10^4^ copies of AST RNA, determined by spectrophotography, were spiked into nucleic acid extracted from 10^5^ uninfected CEM cells (to mimic the 1/1,000 infected cells:uninfected cells in PWH). 2μL of each dilution was added to each PCR reaction well. (**C**) Detection plates of 50μL of select dilutions determined at endpoint. Detection plates were performed with 150μL as well (not shown, but results provided in **Fig. S3D**). (**D**) Average total number of detected AST copies based on the number of positives in the detection plates. Of 10^4^ AST RNA copies, 5,589 were recovered, showing that the sensitivity of our methods for AST detection is about 50%.

**Fig. S4.**
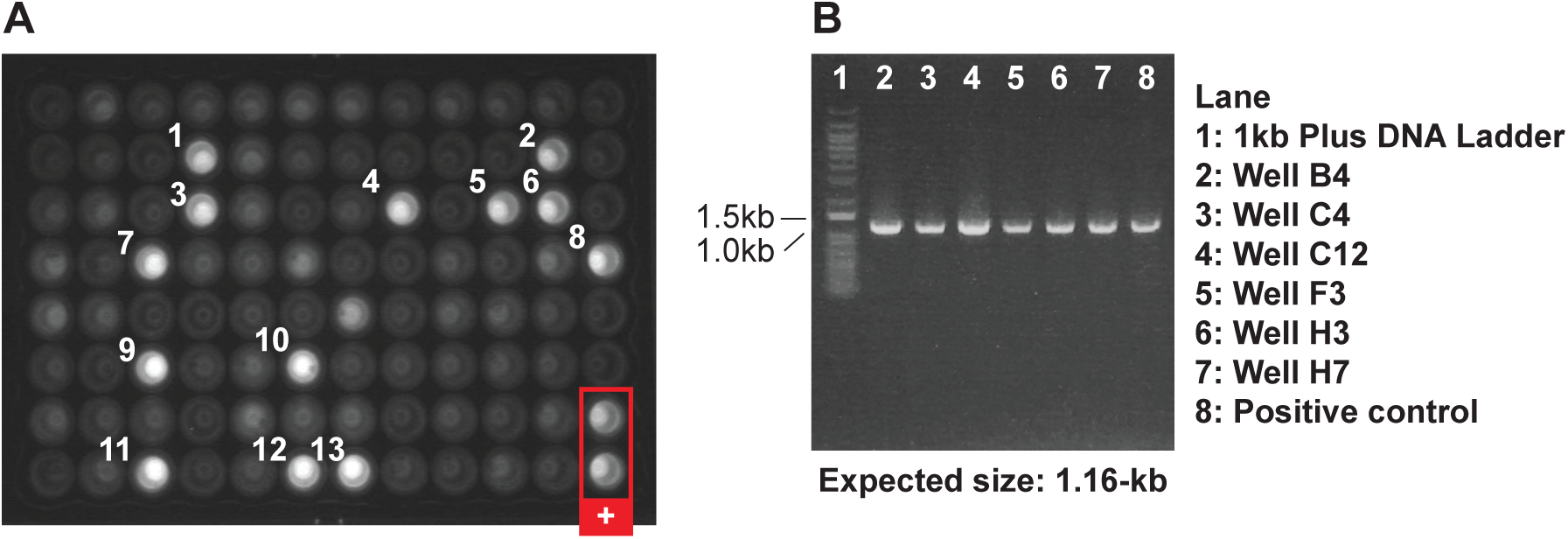
Optimized AST CARD-SGS Assay. (**A**) The *env* AST CARD-SGS assay was applied to an aliquot of 100 ACH-2 cells in a background of 1x10^5^ CEM cells. (**B**) Select positive amplicons from the detection plate in **Fig. S4A** were run on 0.8% agarose gel with the expected amplicon size of ∼1.16-kb.

**Table 2.** Fraction of infected cells with HIV-1 “long AST” in donors on ART.

| Participant Identifier (PID) | Duration on ART at sampling | Estimated number of infected cells assayed | Number of HIV-1 AST sequences obtained | Estimated number of infected cells with HIV-1 AST<sup>A</sup> | Estimate % of infected cells with HIV-1 AST | Average number of HIV-1 AST copies per cell [range]<sup>B</sup> |
| --- | --- | --- | --- | --- | --- | --- |
| 1079 | 12.8 years | 720 | 43 | 12 | 1.7 | 3.6 [1-30] |
| 1683 | 5.4 years | 1,880 | 22 | 20 | 1.1 | 1.1 [1-2] |
| 2669 | 4.3 years | 1,440 | 67 | 63 | 4.4 | 1.1 [1-3] |
|  | 5.5 years | 1,040 | 54 | 52 | 5.0 | 1 [1-2] |
|  | 2 weeks<sup>#</sup> | 2,000 | 86 | 74 | 3.7 | 1.2 [1-3] |
|  | 1 month | 720 | 41 | 40 | 5.6 | 1 [1-2] |
| **Median** |  | **1,240** | **49** | **46** | **4.1** | **1.1** |
| **IQR** |  | **720-1,910** | **36-72** | **18-66** | **1.6-5.2** | **1.0-1.7** |

“long AST” = 1.7kb from 3’ LTR to *env*
<sup>A</sup> Assumes AST with identical sequences are produced in the same single infected cell
<sup>B</sup> Cells without HIV-1 AST were excluded
<sup>#</sup> Not fully suppressed after 2 weeks on ART following 4-week unplanned ART interruption (ATI). PID 2669 was on ART for 5.5 years prior to an ATI. Samples were collected at 4.3 years and 5.5 years prior to the ATI and at 2 weeks and 1 month after ART re-initiation after the ATI

**Table S1.**
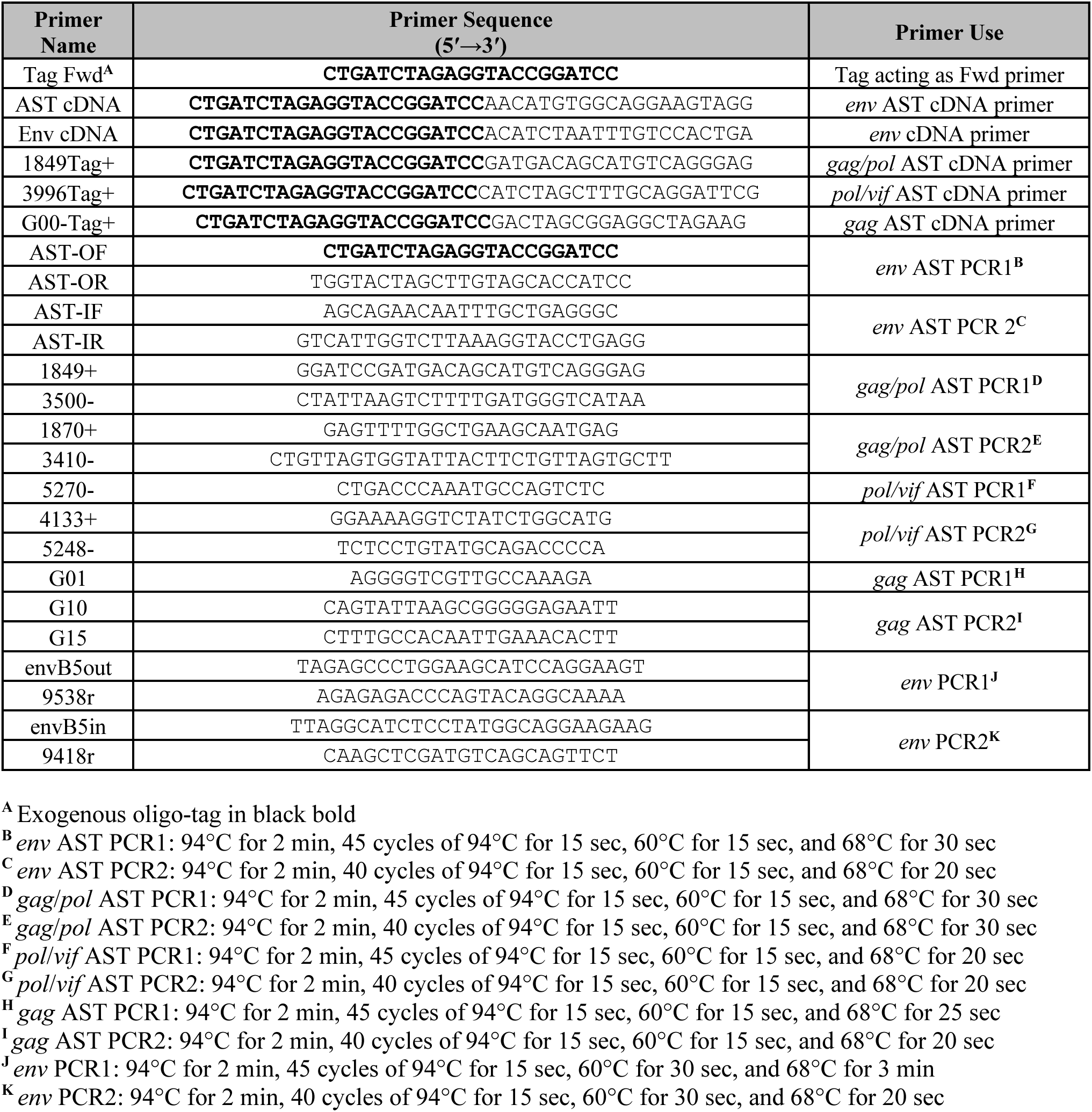
Primers for AST cDNA synthesis and PCR amplification.

**Table S2.**
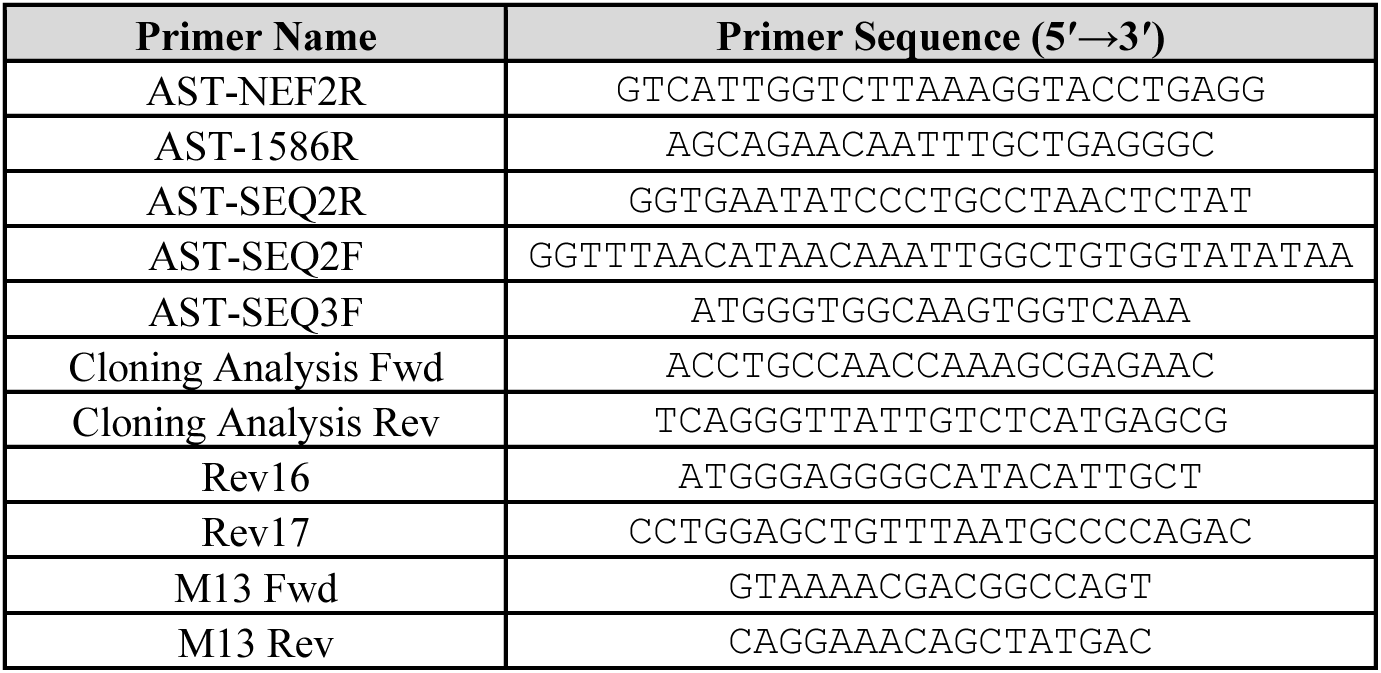
Primers used for AST controls.

**Table S3.**
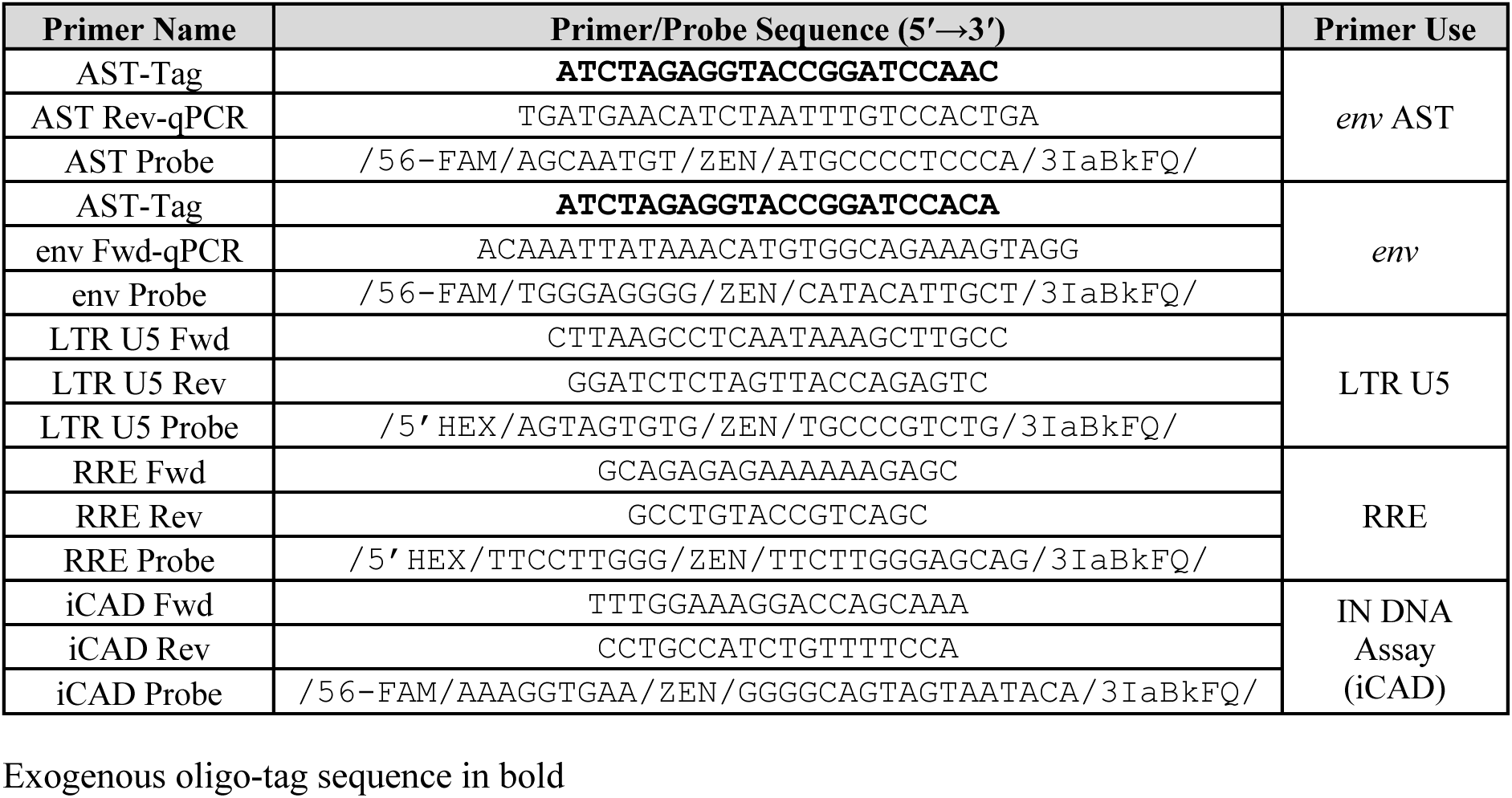
Digital PCR Primers and Probes.

**Table S4.**
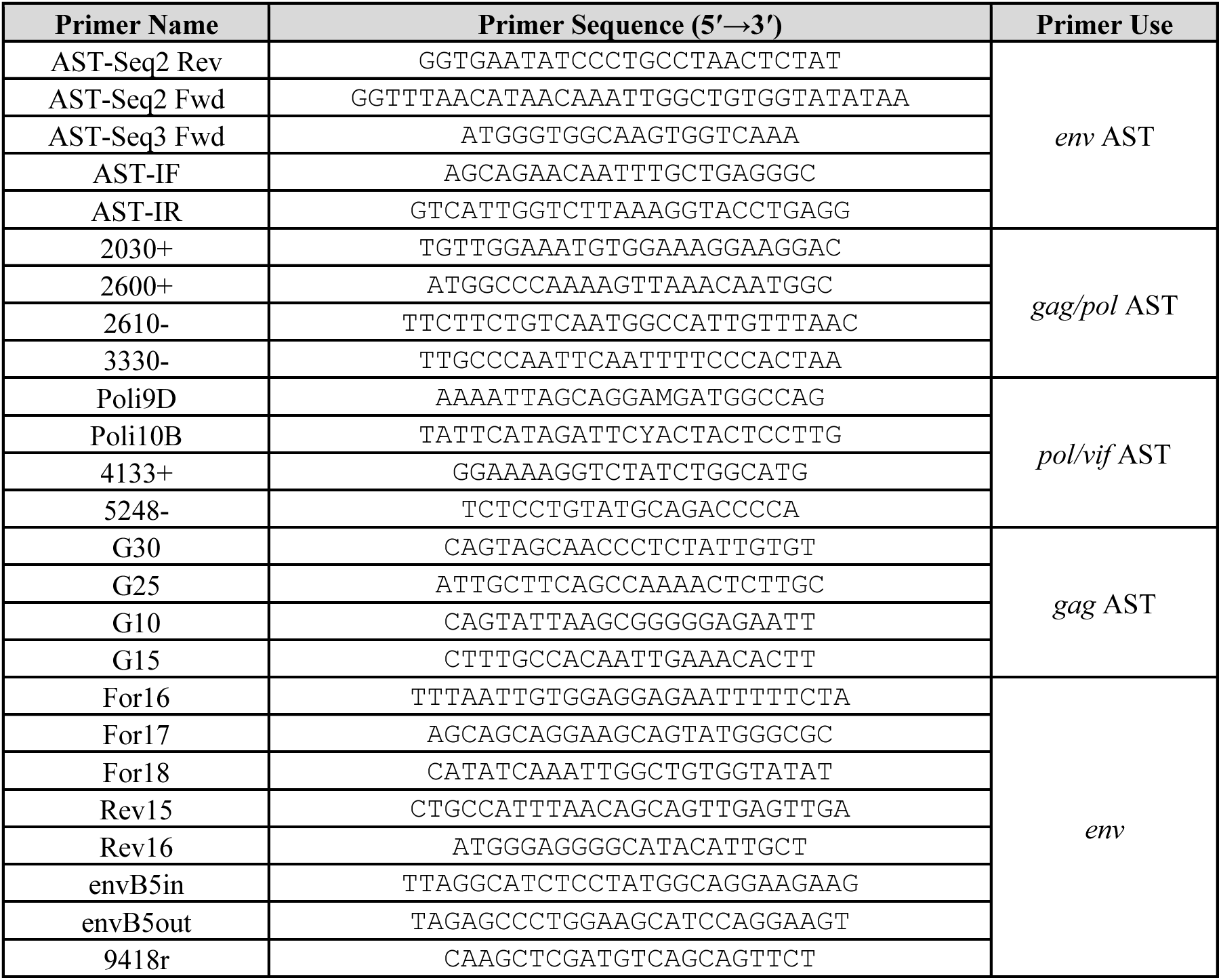
Sequencing Primers.

**Table S5.**
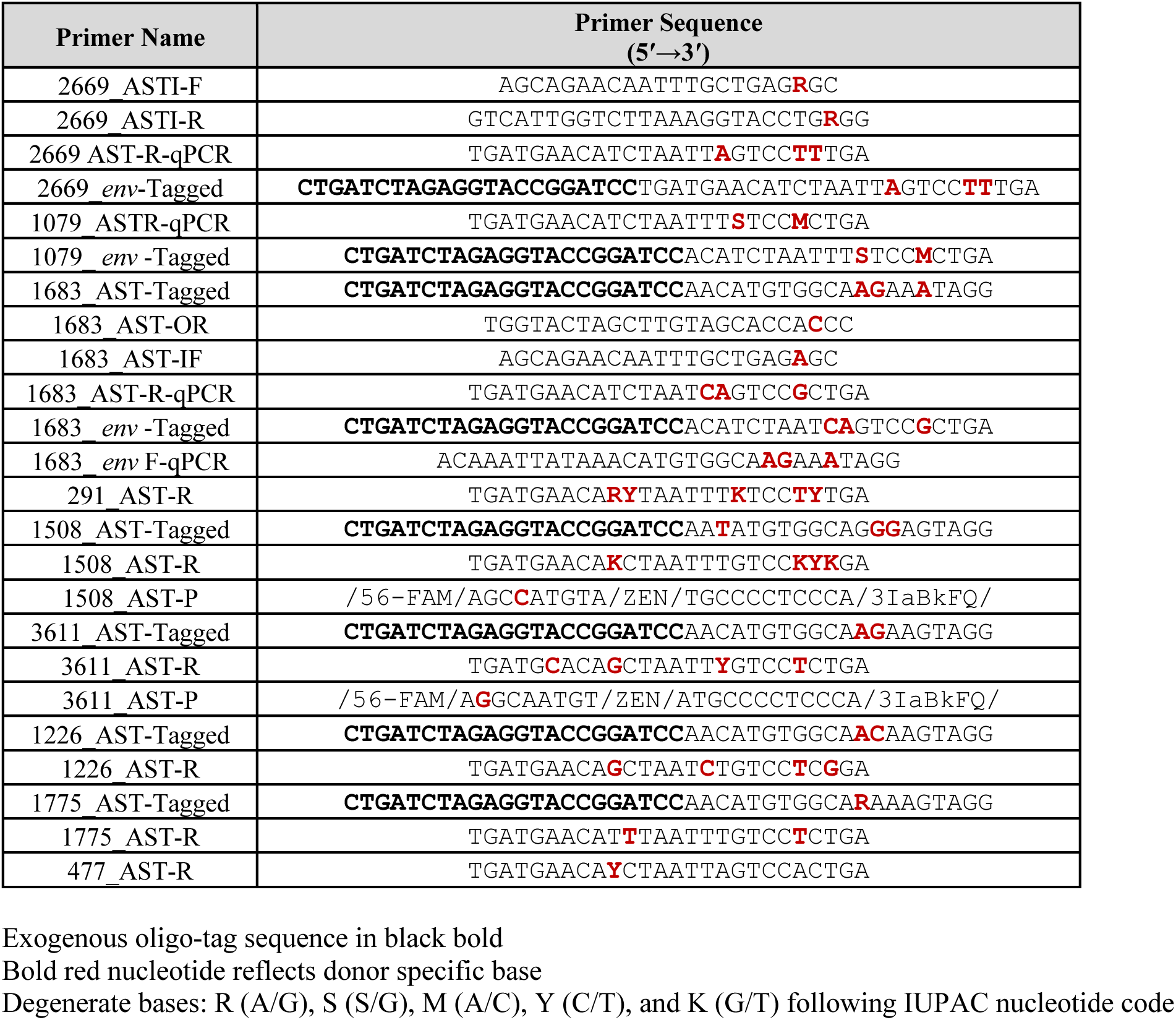
Donor specific primers.

**Table S6.**
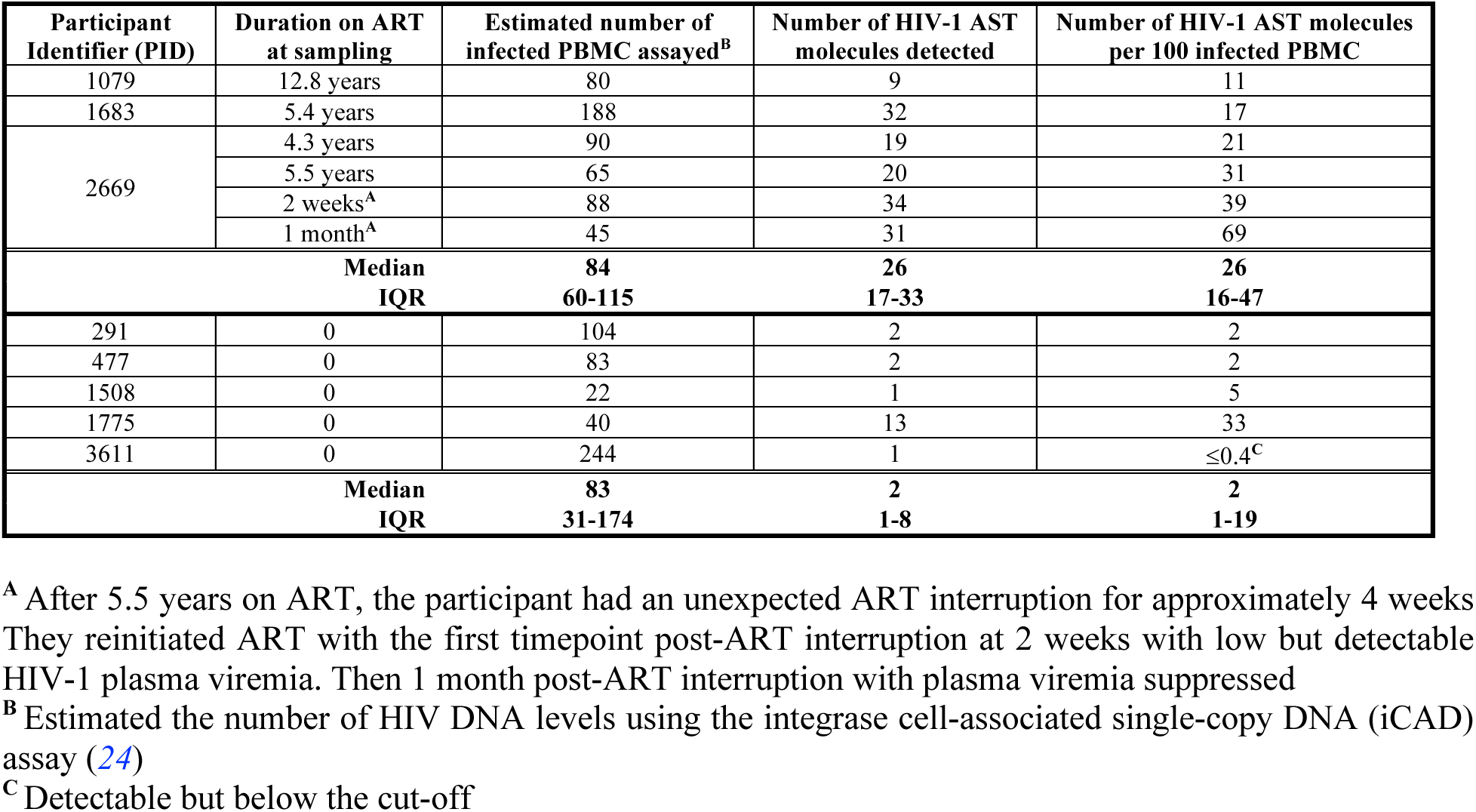
Levels of HIV-1 AST in ART-treated and untreated donors.

## Notes

### Summary of Updates

The revision is to include the NCI funding Project number for this study.

